# Oxygen-enhanced photoacoustic tomography of cerebral hemodynamics in the aging mouse brain, validated against microvascular and microglial histology

**DOI:** 10.64898/2026.11.27.769830

**Authors:** Roshni Parameswaran, B Thamaraiselvi, Ankita Mandal, Thomas Sardella, Sanhita Sinharay

## Abstract

Microvascular rarefaction and impaired cerebral oxygen delivery are hallmark features of the aging brain, linked to age-related cognitive decline and increased stroke susceptibility. Yet noninvasive methods to track them in preclinical models remain limited. We used photoacoustic tomography, which combines label-free molecular contrast from endogenous hemoglobin in its oxygenated and deoxygenated forms with ultrasound-defined spatial resolution, to obtain surrogate markers of brain hemodynamics. Using oxygen-enhanced multispectral optoacoustic tomography (OE-MSOT) with non-negative model-based (NNMB) reconstruction, we measured MSOT-derived oxy- and deoxyhemoglobin (HbO₂ and Hb), total hemoglobin (HbT) and oxygen saturation (sO₂^MSOT^) in a whole-slice brain region of interest in mice aged 3, 6, 12 and 15 months, under normoxia (21% O₂) and hyperoxia (100% O₂). We compared imaging readouts with CD31 (microvascular) and IBA1 (microglial) immunofluorescence in the same animals at 3 and 15 months. Under normoxia, 15-month-old mice showed lower HbO₂ (−33%) and HbT (−26%) and a 4.1 percentage point lower baseline sO₂^MSOT^ than 3-month-old mice, accompanied by a 56% reduction in CD31⁺ vascular density and reduced microglial ramification (animal-level analysis, p ≤ 0.04). Vascular density correlated strongly with HbO₂ and HbT across animals (Spearman ρ = 0.90-0.92, p < 0.001), with a consistent positive relationship at both ages. Hyperoxia increased sO₂^MSOT^ at every age, and the magnitude of this response (ΔsO₂^MSOT^) declined with age, from a mean of 5.5% at 3-months to 3.3% at 15 months (linear regression, p = 0.034). These findings indicate that OE-MSOT blood-volume readouts track histological microvascular rarefaction in the aging mouse brain, supporting their use as noninvasive, longitudinally compatible markers of cerebrovascular aging.

## 1. Introduction

Structural and functional changes in cerebral vasculature are concurrent with aging and lead to hypoperfusion and cognitive decline, lowering the threshold for neurodegenerative disease(1–3). Distinguishing normal age-related changes from the progressive, pathological neuronal loss of neurodegenerative disease is therefore an important goal. Age-related changes in vessel morphology and the associated hemodynamic variations are implicated in accelerated cognitive decline in the elderly(4–6) underscoring the need for tools that can track cerebral oxygenation across the lifespan. Mouse models are powerfully suited to address this question: they recapitulate key features of human cerebrovascular aging, permit genetic manipulation and allow longitudinal assessment within a 2-3-year lifespan(6–8).

Magnetic resonance imaging (MRI) has characterized the aging brain with manifestations of brain atrophy, white-matter hyperintensities, progressive accumulation of clinically silent microbleeds and small infarcts, indicative of compromised vascular integrity in the aged brain(9–11). These anatomical features, together with altered metabolism and a low-grade neuroinflammatory state, capture well the landscape of the aging brain(12–14). Alterations in vascular morphology and blood flow dynamics parallel these structural and metabolic changes(15). In humans, transcranial and carotid Doppler ultrasonography shows an age-associated decrease in blood flow velocity in large cerebral arteries(16, 17), and magnetic resonance perfusion imaging confirms a global decline in cerebral blood flow (CBF) with age(11, 15). At the medium-vessel level, MR angiography reveals fewer visible cerebral vessels with age in humans, and aged mice demonstrate fewer and more tortuous cerebral arterioles(18–20). At the microvascular level, both species show age-related reductions in capillary density (21, 22). In parallel, microglia, the resident immune cells of the brain and a constituent of the neurovascular unit, also undergo morphological remodeling with age that differs from the reactive transformation observed in acute neuroinflammation or neurodegenerative diseases and may interact with vascular changes to collectively shape the aged neurovascular environment(23).

Whether these structural changes cause reduced perfusion, impaired oxygen delivery, and neuronal dysfunction remains incompletely understood. Resolving this question requires noninvasive functional imaging of cerebral oxygenation that can be performed longitudinally and compared directly against structural readouts. Clinically, cerebral perfusion and oxygen metabolism are measured with positron emission tomography (PET), using [^15^O] H_2_O for CBF, [^15^O] O_2_ for oxygen extraction fraction and metabolic rate, and ^18^F-FDG for glucose metabolism(24, 25). Translating these measurements to mice is difficult: [^15^O]-labeled tracers require an on-site cyclotron (∼2 min half-life), and small-animal PET has limited spatial resolution relative to the size of the mouse brain, with repeated studies constrained by the radiation dose, anesthesia, and cost. Blood Oxygen Level Dependent MRI (BOLD MRI) provides oxygenation-sensitive contrast with good spatial coverage, but the detected signal reflects combined changes in blood flow, blood volume and oxygen consumption, rather than direct measurements of oxygen saturation, and preclinical high-field MRI is costly as well as technically demanding, making its routine use challenging(26, 27). Near-infrared spectroscopy (NIRS) is portable and sensitive to oxygenation but limited in spatial resolution and depth (28–30).

These modalities are increasingly combined with gas challenges to prove vascular function (31). Hypercapnia, used to measure cerebrovascular reactivity (CVR), evaluates the capacity of cerebral vessels to dilate in response to a vasoactive stimulus and has emerged as a clinically validated biomarker of vascular contributions to cognitive impairment(32, 33). Hyperoxia, in contrast, is not a vasodilatory stimulus and causes mild cerebral vasoconstriction(34). The resulting rise in tissue oxygenation therefore reflects the combined effect of increased arterial oxygen content and hyperoxic vasoconstriction on oxygen delivery through existing vasculature. Oxygen-enhanced MRI and photoacoustic imaging have used this response as a surrogate of vascular function(35, 36).

Photoacoustic (optoacoustic) imaging combines optical absorption contrast with ultrasonic detection and offers a trade-off between resolution and depth across its implementations. Photoacoustic microscopy (PAM) resolves individual microvessels near the brain surface and has been used to quantify hemoglobin concentration, oxygen saturation, and blood flow through the intact mouse skull(37) and to characterize disease-related microvascular changes in the brain(38). Photoacoustic tomography (PAT) trades this resolution for depth, providing sub-millimeter resolution at depths of several centimeters, and has been used to map cerebral hemodynamics during sensory stimulation(39) and in glioblastoma(40). Multispectral optoacoustic tomography (MSOT) acquires images at several wavelengths to delineate signatures of endogenous oxyhemoglobin (HbO₂) and deoxyhemoglobin (Hb), giving label-free molecular contrast for blood volume and oxygen saturation (sO₂^MSOT^), where the reliability of these readouts depends on how the images are reconstructed. Model-based reconstruction with a non-negativity constraint (NNMB) reduces the negative-value artifacts of standard back-projection (BP) and improves the reliability of spectral unmixing, although the resulting parameters remain rather relative than absolute measures(41, 42). Combined with a hyperoxic challenge, oxygen-enhanced MSOT (OE-MSOT) has been validated against histology as a surrogate of vascular function and hypoxia in tumors(43), but to our knowledge it has not been applied systematically to the aging brain or compared with histology in the same animals. Clinical MSOT systems are already used to assess hemoglobin, oxygenation and other endogenous chromophores as imaging biomarkers in inflammatory, muscular diseases and in cancer(44, 45). Establishing what MSOT-derived cerebral readouts reflect at the tissue level is therefore also a step towards their use as translational biomarkers.

In this study, we used OE-MSOT with NNMB reconstruction to measure cerebral hemodynamics in female BALB/c mice at 3, 6, 12, and 15 months of age, under normoxia (21% O_2_) and hyperoxia (100% O_2_), in a whole-slice region of interest (ROI) at a single coronal plane. To relate the imaging readouts to their structural and cellular basis, we performed immunofluorescence on the brains of imaged 3-month and 15-month-old mice, staining the microvasculature with the endothelial marker CD31 and microglia with IBA1, and compared imaging and histology in the same animals. Endothelial GLUT1 was assessed qualitatively. We hypothesized that aging would reduce MSOT-derived blood volume and baseline sO₂^MSOT^ and attenuate the oxygen-challenge response (ΔsO₂^MSOT^), and that these changes would track microvascular rarefaction, providing a noninvasive readout of the aged neurovascular unit.

## 2. Methods

### 2.1 Animals and study design

All experimental protocols in this study were approved and performed in accordance with the Control and Supervision Rules, 1998 of the Ministry of Environment and Forest Act (Government of India), and the Institutional Animal Ethics Committee, Indian Institute of Science (protocol no. IISc CAF/ethics/099/2024). Mice were procured from the Central Animal Facility (CAF) of the Indian Institute of Science (IISc, Bangalore). All experiments were performed in female BALB/c strains of mice. We examined age-group differences in a cross-sectional design with four age groups: n=8 (3 months-old), n=5 (6 months-old), n=5 (12 months-old), and n=7 (15 months-old). The ages were chosen to span young adulthood to middle age in mice, consistent with previous studies of cerebrovascular aging(46). Due to technical challenges with anesthesia during imaging, the MSOT data of one 12-month-old and one 15-month-old were excluded from all analyses, bringing the total to n=23. Post-imaging, mice were perfused and the brains of 3-month-old and 15-month-old mice were collected for immunofluorescence studies.

### 2.2 Magnetic resonance imaging

To localize the photoacoustic imaging plane anatomically, one 3-month-old mouse underwent both MRI and MSOT. MRI was performed on a 3T scanner (Prisma, Siemens Healthineers) with a 32 mm inner-diameter ¹H volume coil (RAPID Biomedical, Germany). The mouse was anesthetized (ketamine+xylazine) and the head was positioned at the coil isocenter and coronal T2-weighted turbo spin-echo (TSE) images were acquired (TR/TE = 4620/78 ms; field of view 45 × 85 mm; matrix 170 × 320; 20 slices; slice thickness 1 mm; acquisition time 8 min). MSOT and MR slices were matched using the lateral ventricles and prominent vessels as landmarks, and the imaging plane was assigned to an atlas level using the anterior commissure crossing the midline (MR slice 7, **Fig.1A, B**) and the first appearance of the dorsal hippocampus (MR slice 8, Fig.1), corresponding to approximately Bregma −0.10 to −1.22 mm(47).

**Figure 1.**
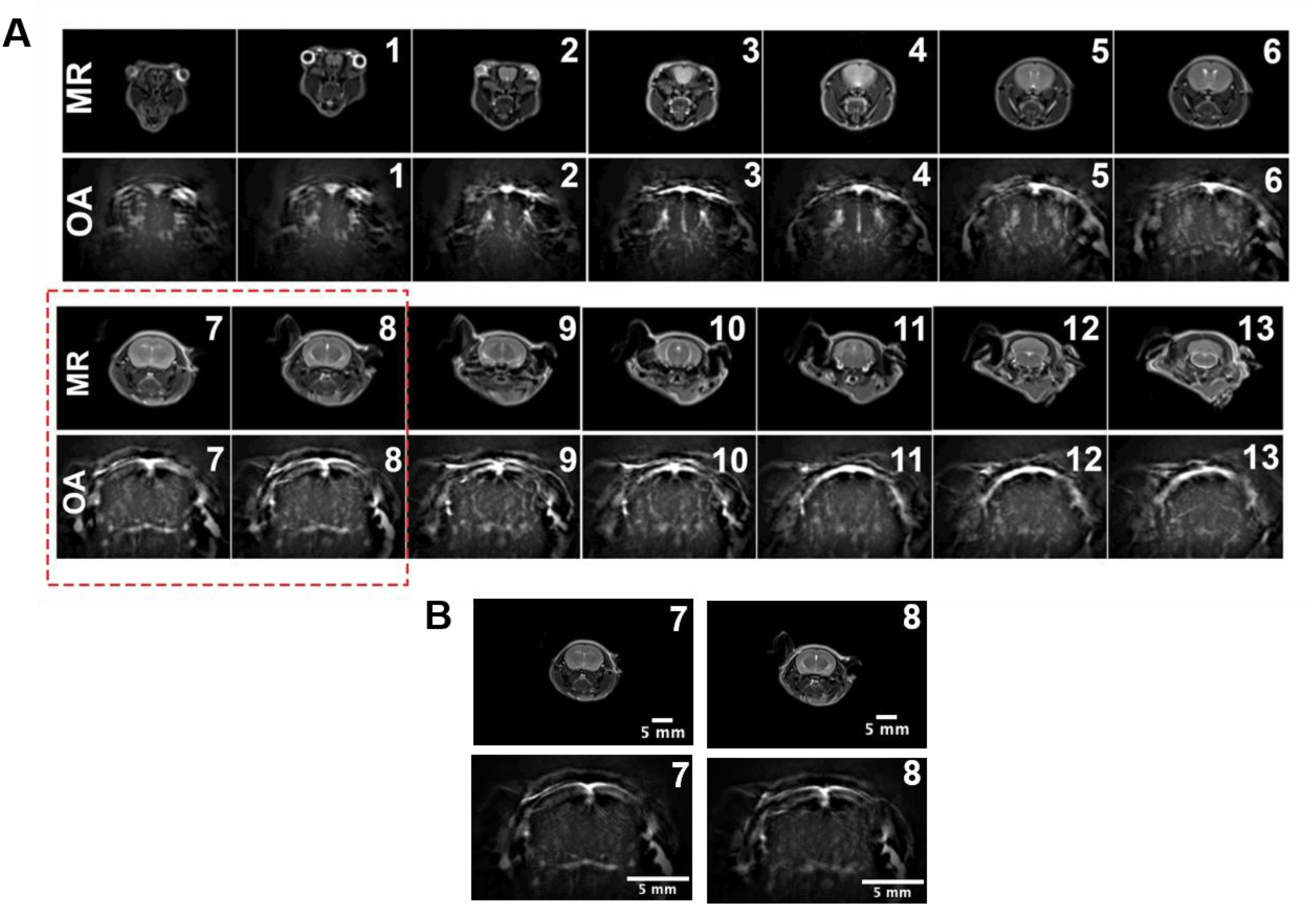
Anatomical localization of the MSOT imaging plane. (A) Paired coronal T2-weighted MR and single-wavelength (800 nm) optoacoustic (OA) images from the same 3-month-old mouse, from slice 1 (olfactory bulb) to slice 13 (cerebellum and brainstem). MR slices are 1 mm thick and OA slices 0.5 mm; OA slices shown are those that most closely matched each MR slice. The red dashed box marks slices 7 and 8, which contain the anterior commissure crossing the midline and the first appearance of the dorsal hippocampus. (B) Enlarged MR (top) and OA (bottom) images of slices 7 and 8. Scale bars, 5 mm

### 2.3 Oxygen-enhanced MSOT imaging

For brain MSOT imaging, mice were prepared by shaving the head and positioned supine in the holder. Mice were induced under 3 % isoflurane mixed with 100% O_2,_ flow rate of 1L/min. Mice were secured in a customized animal holder (iThera Medical, Germany) with a bite bar, and limbs secured through elastic handcuffs to maintain a taut position with minimal movement. The physiological body temperature was maintained by keeping the water bath in the MSOT scanner at 36 ℃ during the scans. Respiratory rate was monitored visually and kept at approximately 60 breaths/min.

Imaging was performed on an MSOT inVision 256-TF system (iThera Medical, Germany) using nanosecond laser pulses at 10 Hz and a 256-element ring transducer array (5 MHz centre frequency). MSOT signals were corrected for pulse-to-pulse variation in laser energy using the system’s built-in per-pulse energy normalization(48, 49). Post-induction, the anesthesia was kept at 2% for subsequent scans. For imaging, the breathing air was first switched to medical air (21% O₂) for 8 minutes, following which a multi-slice scan covering the brain and skull (a total of 29 positions) was acquired at wavelengths 700, 730, 760, 800, 850 and 875 nm; 0.5 mm step interval; 10 frames averaged per wavelength; and a total scan time of 5 min. From this scan, the analysis plane was selected in each mouse as described in the previous section, corresponding to approximately Bregma −0.10 to −1.22 mm. A single-slice acquisition at this plane was then performed (same six wavelengths; 10 frames per wavelength; 50 repetitions; 5 min). The inspired gas was then switched to 100% O₂ and, after a further 8-minute equilibration, the single-slice acquisition was repeated with identical settings. The 8-minute equilibration was chosen to allow tissue oxygenation to reach a steady state after each gas change(43, 50). Further, to illustrate the time course of the oxygen-challenge response, a dynamic acquisition was performed in one 3-month-old mouse, imaging the analysis plane continuously (6 wavelengths, 10 frames per wavelength) while the inspired gas was switched from 21% to 100% O₂) (**Fig. 2C, D, E**). The gas order was the same for all animals (21% followed by 100% O₂), and the total anesthesia time was approximately 30 min per mouse after induction.

**Figure 2.**
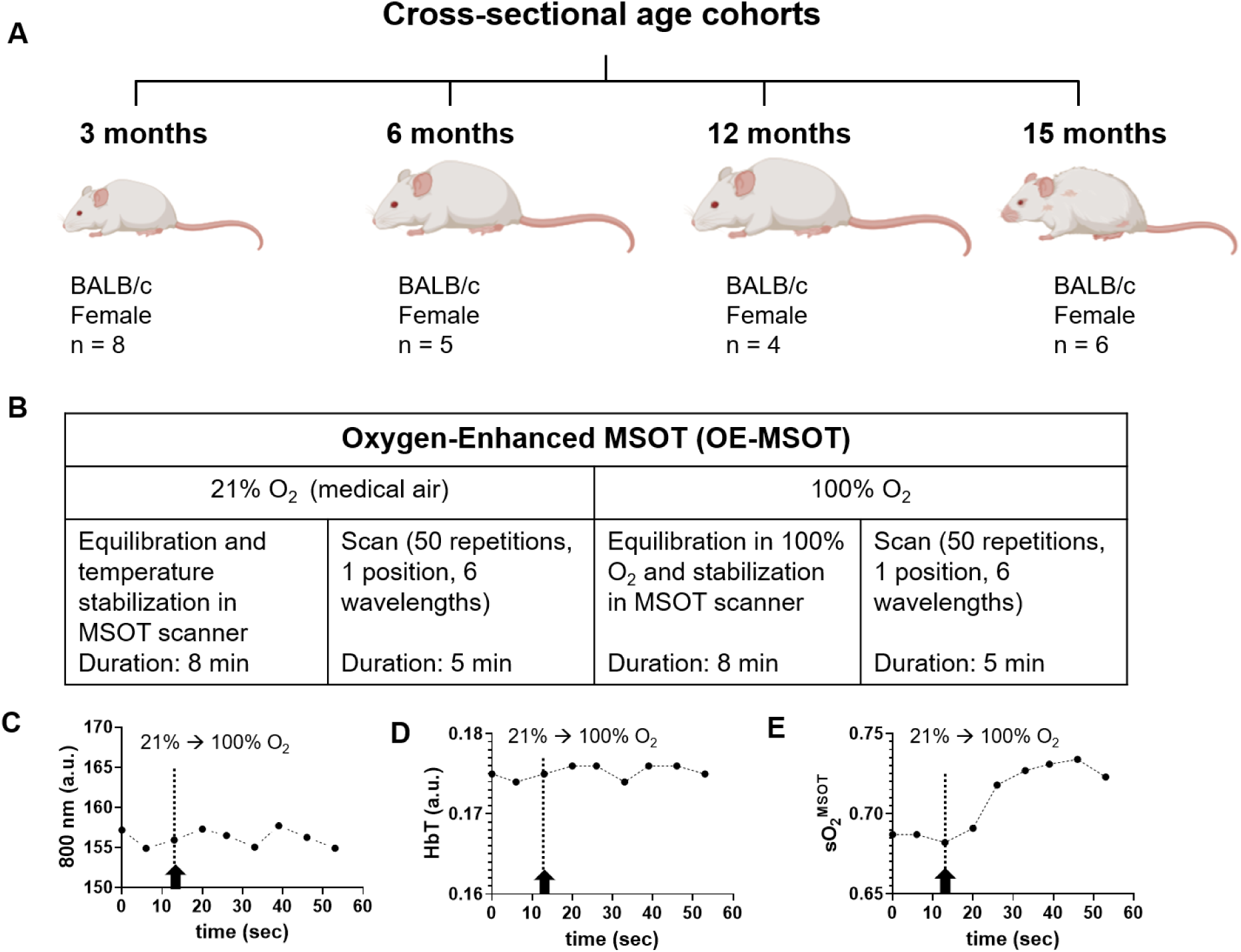
Study design and oxygen-enhanced MSOT imaging paradigm. (A) Cross-sectional design with four age groups of female BALB/c mice: 3 months (n = 8), 6 months (n = 5), 12 months (n = 4) and 15 months (n = 6). (B) Single-slice OE-MSOT acquisition at the analysis plane. (C) Representative dynamic acquisition during the gas switch (arrow and dashed line, 21% → 100% O₂) in one 3-month-old mouse, showing the photoacoustic signal at 800 nm (left), (D) HbT (middle) and (E) sO₂^MSOT^ (right) within the whole-slice ROI.

### 2.4 Image reconstruction and analysis

Images were reconstructed with a non-negative constrained model-based algorithm (NNMB; iLabs v1.3.33, iThera Medical) (41, 42) using a 25 mm field of view, 100 µm pixel size, speed of sound 1530 m/s and a 50 kHz–6 MHz band-pass filter. For comparison, images were also reconstructed by back-projection (BP) in viewMSOT 4.0 (keeping the same field of view, speed of sound, and filter settings). NNMB reconstructions were imported into viewMSOT 4.0 for further processing, including fluence correction and spectral unmixing.

A mask covering the brain cross-section was manually drawn on each image and used for fluence correction with the built-in viewMSOT model. The tissue optical properties are fixed in the software (water volume fraction 0.16, fat volume fraction 0.62, scattering decay 1.05, scattering scaling 16.7 cm⁻¹), and the blood volume fraction was left at its default of 0.0098. Spectra were unmixed pixel-wise by linear regression against the absorption spectra of Hb and HbO₂, yielding maps of Hb and HbO₂. HbT was calculated pixel-wise as Hb + HbO₂, and %sO₂MSOT as HbO₂/HbT × 100. The values for each parameter were averaged within a whole-slice ROI covering the entire brain cross-section, excluding scalp and skull, and then across the 50 repetitions. Because fluence correction is approximate, Hb, HbO₂, and HbT are reported as relative values in arbitrary units. The oxygen-challenge response was calculated per animal as ΔsO₂MSOT = %sO₂MSOT (100% O₂) − %sO₂MSOT (21% O₂). Changes in sO₂MSOT are reported as absolute differences in % sO₂; changes in Hb, HbO₂, and HbT are reported as percentage change relative to the 3-month group. To assess the contribution of superficial tissue, an ROI was manually drawn along the scalp overlying the dorsal brain in the same imaging plane under 21% O₂ (**Fig. S2A, B**). The mean photoacoustic signal at 800 nm (before spectral unmixing) and mean HbT within this ROI were averaged across the 50 repetitions and compared between 3- and 15-month-old mice. mice.

### 2.5 Tissue processing

After imaging, mice were anesthetized with isoflurane and transcardially perfused with 1X phosphate-buffered saline (PBS), followed by 4% paraformaldehyde (PFA). After perfusion, brains were carefully dissected and post-fixed in 4% PFA for 24 hours. The fixed tissues were then cryoprotected in 30% sucrose prepared in 1X PBS (w/v) for 24-48 hours, embedded in optimal cutting temperature (OCT) compound, and coronally sectioned at a thickness of 50 µm using a sliding cryotome (RWD. Minux, FS800A) maintained at −20 ℃. Sections at approximately Bregma 0 to −1 mm, matching the MSOT imaging plane, were used for staining (**Fig. S5**).

### 2.6 Immunofluorescence and image quantification

Free-floating sections were incubated overnight at 4 °C with primary antibodies against CD31 (rat anti-mouse, 1:100, BD Pharmingen), IBA1 (rabbit monoclonal, 1:250, Thermo Fisher Scientific) or GLUT1 (rabbit monoclonal, 1:100, Cell Signaling Technology), followed by Alexa Fluor 488-conjugated anti-rat (1:300) or anti-rabbit (1:500) or Alexa 594-conjugated anti-rabbit (1:350) secondary antibodies, and DAPI. CD31, IBA1 and GLUT1 were stained on separate sections. Secondary-only controls were included; and autofluorescence was assessed in unstained sections imaged with identical settings.

Z-stacks were acquired with a 60× (or 40×) objective on a Leica SP8 (CD31) or Olympus FV3000 (IBA1, GLUT1) confocal microscope, using identical acquisition settings for both age groups, and displayed as maximum-intensity projections of 50 µm. Images were analyzed in Fiji/ImageJ. For CD31, vascular density and vascular length density were measured with the Vessel Analysis plugin using Li thresholding algorithm, in 10 fields (184 × 184 µm²) from the cortex region per mouse, averaged to one value per animal. For IBA1, individual microglia fields (10-15 cells per mouse) were converted to 8-bit images, processed using threshold algorithm, skeletonized, and analyzed with AnalyzeSkeleton to obtain the number of branches, junctions, and endpoints and the maximum branch length per cell, averaged per animal. Histology was quantified in imaged 3-month-old mice with available tissue (n=6) and all 15-month-old mice in the final cohort (n=7). GLUT1 was assessed qualitatively in two mice per age group.

### 2.7 Statistical analysis

All statistical analyses were performed in GraphPad Prism (version 10.6, GraphPad Software, USA) and effect sizes in Microsoft Excel. Data are presented as mean ± SD with individual animals shown. Differences across the four age groups in Hb, HbO₂, HbT and sO₂^MSOT^, under 21% and 100% O₂, were tested by one-way ANOVA followed by Dunnett’s multiple comparisons test against the 3-month group. The change in %sO₂^MSOT^ between 21% and 100% O₂ within each age group was tested by two-tailed paired t-test. The relationship between ΔsO₂^MSOT^ and age was assessed by linear regression with age (months) as a continuous predictor.

Comparisons between 3- and 15-month-old mice (sO₂^MSOT^ with NNMB and BP reconstruction, and all histological measures) used two-tailed unpaired t-tests with Welch’s correction. Effect sizes are reported as Hedges’ g (Cohen’s d with small-sample correction) with approximate 95% confidence intervals. Whole-slice ROI area was compared across age groups by one-way ANOVA, and scalp signal between 3 and 15 months by unpaired t-test.

Pooled associations were estimated by Spearman correlation (Python). To separate the imaging-histology relationship from the shared effect of age, associations within each age group were assessed by linear regression and the slopes were compared between age groups and the age-adjusted associations were tested by analysis of covariance (ANCOVA), with the MSOT readout as the dependent variable, age group as a factor and CD31⁺ vascular density as the covariate, after confirming homogeneity of slopes between age groups. The relationship between MSOT HbO₂/HbT and CD31⁺ vascular density was specified in advance as the primary association of interest; and other imaging histology associations were treated as exploratory. Correlations among imaging parameters were not interpreted, because several are mathematically related (HbT=Hb + HbO₂; ΔsO₂^MSOT^ is derived from sO₂^MSOT^).

A two-sided p < 0.05 was considered significant. In figures, * p < 0.05, ** p < 0.01, *** p < 0.001, **** p < 0.0001

## 3. Results

### 3.1 Anatomical localization of the MSOT imaging plane

To define the anatomical position of the coronal plane used for hemodynamic analysis, we acquired MSOT images across the whole brain (800 nm, 0.5 mm slice thickness) and compared them with the T2-weighted MR images (51, 52) acquired in the same 3-month-old mouse (**Fig. 1**), on a 3T MRI (Siemens Prisma, equipped with a preclinical volume coil (Inner diameter = 32 mm, RAPID Biomedical, Germany). The MSOT series reproduced the brain outline, bilateral symmetry and anterior-to-posterior progression seen in the MR series, although with lower soft-tissue contrast. The lateral ventricles and the prominent surface vessels served as the landmarks for matching the sections across the two modalities. The closest correspondence in hemisphere shape and parenchymal boundaries was found for MSOT slices 7-8 (**Fig. 1**, red dashed box), corresponding to Bregma 0 to −1 mm in the mouse brain. At this level, the brain cross-section comprises the cortex, corpus callosum and caudate-putamen, together with the rostral dorsal hippocampus and thalamus at its caudal end. In all subsequent animals, the analysis plane was identified in the multislice MSOT scan using the same landmarks, and hemodynamic parameters were measured in a whole-slice ROI covering the entire brain cross-section at this level.

### 3.2 NNMB reconstruction preserves the dynamic range of sO₂^MSOT^ and resolves age-group differences

Cerebral hemodynamics were assessed in a cross-sectional cohort of female BALB/c mice aged 3, 6, 12 and 15 months. Of the 25 mice imaged, two (one each at 12 and 15 months) were excluded due to technical issues, giving final imaging group sizes of n=8, 5, 4 and 6, respectively. Each mouse was imaged in a single session under medical air (21% O₂) followed by 100% O₂, with an 8-minute equilibration period before each acquisition (**Fig. 2A, B**) (40). The change in sO₂^MSOT^ between the two conditions (ΔsO₂^MSOT^) was evaluated as the oxygen-challenge response. A dynamic acquisition during the gas switch in a representative 3-month-old mouse showed that sO₂^MSOT^ began to rise within seconds of switching to 100% O₂, while HbT and the 800 nm signal remained stable (**Fig. 2C-E; Fig. S1**). The change in oxygenation therefore occurred without a detectable change in blood volume or overall signal intensity and developed well within the 8-minute equilibration period.

Images were reconstructed with a non-negative constrained model-based algorithm (NNMB)(41, 42) followed by linear unmixing of the Hb and HbO₂ spectra. Standard back-projection (BP) produces negative pixel values with no physical meaning that propagate into the unmixed maps; whereas constraining reconstructed values to be non-negative through NNMB reconstruction removes these artifacts, as seen in the sO₂^MSOT^ maps (**Fig. 3A, B**). To assess the effect of this choice, sO₂^MSOT^ was compared between 3- and 15-month-old mice using both reconstructions. sO₂^MSOT^ was lower in 15-month-old mice with both methods, but the difference was larger and statistically significant with NNMB (21% O_2_: NNMB, Hedges’ g=2.0, 95 % CI 0.7-3.3, p=0.007; BP, g=0.8, 95% CI −0.3-1.9, p=0.16; 100% O₂: NNMB, g=2.3, 95% CI 1.0-3.7, p < 0.001; BP, g=1.0, 95% CI −0.2 to 2.1, p=0.11) (**Fig. 3C, D**). BP compressed the overall range of sO₂^MSOT^ values where the response to hyperoxia was approximately three-fold smaller with BP than with NNMB in 3-month-old mice, consistent with the negative-value artefacts biasing the spectral unmixing. NNMB reconstructions were used for all subsequent analyses.

**Figure 3.**
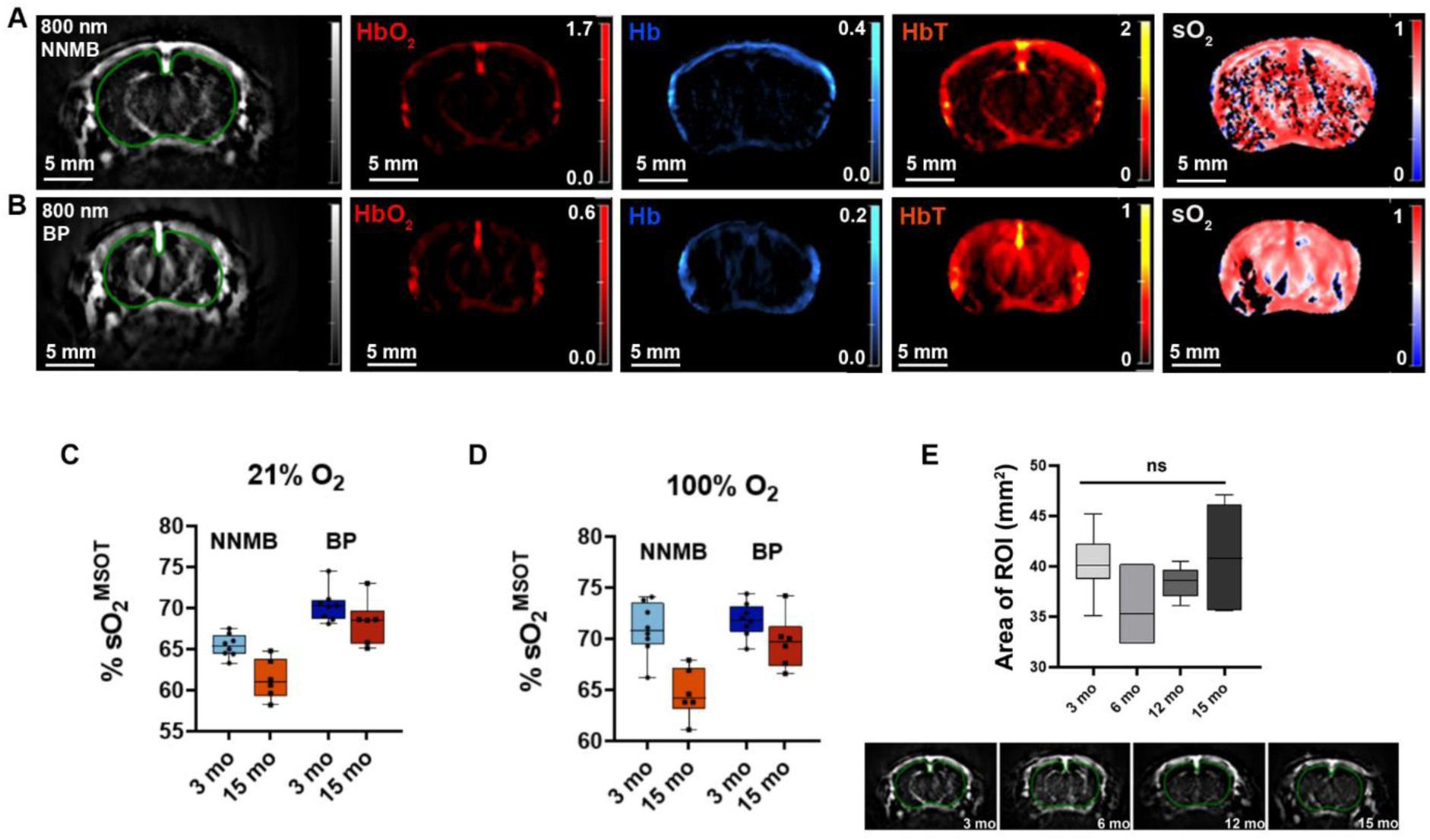
Comparison of reconstruction algorithms and the size of whole-slice ROI. (A, B) Representative 800 nm images and HbO₂, Hb, HbT and sO₂^MSOT^ maps of a 3-month-old mouse brain reconstructed with (A) non-negative model-based (NNMB) and (B) back-projection (BP) algorithms (100 µm pixel size). The green outline depicts the whole-slice ROI. (C, D) Whole-slice sO₂^MSOT^ in 3-month-old (n = 8) and 15-month-old (n = 6) mice reconstructed with NNMB and BP under (C) 21% O₂ and (D) 100% O₂; each point represents one mouse. (E) Whole-slice ROI area across age groups (3 months, n = 8; 6 months, n = 5; 12 months, n = 4; 15 months, n = 6; one-way ANOVA, p = 0.09), with representative ROIs for each age shown below (green trace).

Three control analyses were performed. First, whole-slice ROI area was compared between the four groups and did not differ significantly between age groups (one-way ANOVA, p=0.09; **Fig. 3E**), indicating that the group differences were not driven by whole-slice ROI size. Second, to assess the contribution of superficial tissue, the photoacoustic signal at 800 nm and HbT were measured in a scalp ROI overlying the brain (**Fig. S2A, B**). Neither differed between 3- and 15-month-old mice (unpaired two-tailed t-test, p=0.52 and p=0.68; **Fig. S2C, D**), indicating comparable superficial signal at both ages. This analysis does not, however, exclude age-related differences in light and sound attenuation by the skull itself, which are considered in the discussion later. Third, to examine whether whole-slice values were representative of the underlying tissue, %sO₂^MSOT^ was measured in regional sub-ROIs of the imaging plane (**Fig. S3A**). In a representative3-month-old mouse, cortical and striatal %sO₂^MSOT^ were similar to the whole-slice value under both 21% and 100% O₂, and %sO₂^MSOT^ increased in all regions under hyperoxia including the midline and lateral surface vessels (**Fig. S3B**).

### 3.3 Aged mice show lower MSOT-derived blood volume and oxygen saturation

We next evaluated MSOT-derived Hb, HbO_2_, HbT and %sO₂^MSOT^ across the four age groups under 21% O_2_ followed by 100% O_2_ in inspired air (**Fig. 4A-H**). Within the 680-900 nm wavelength range oxy- and deoxyhemoglobin dominate optical absorption in brain tissue, and other chromophores such as water and lipids absorb at least an order of magnitude less (53). Fluence-corrected photoacoustic measurements within the chosen ROI provided quantitation of Hb and HbO_2_ and their sum (Hb+HbO_2_), represented HbT, which is another metric for regional cerebral blood volume (CBV) within the tissue(54, 55).

**Figure 4.**
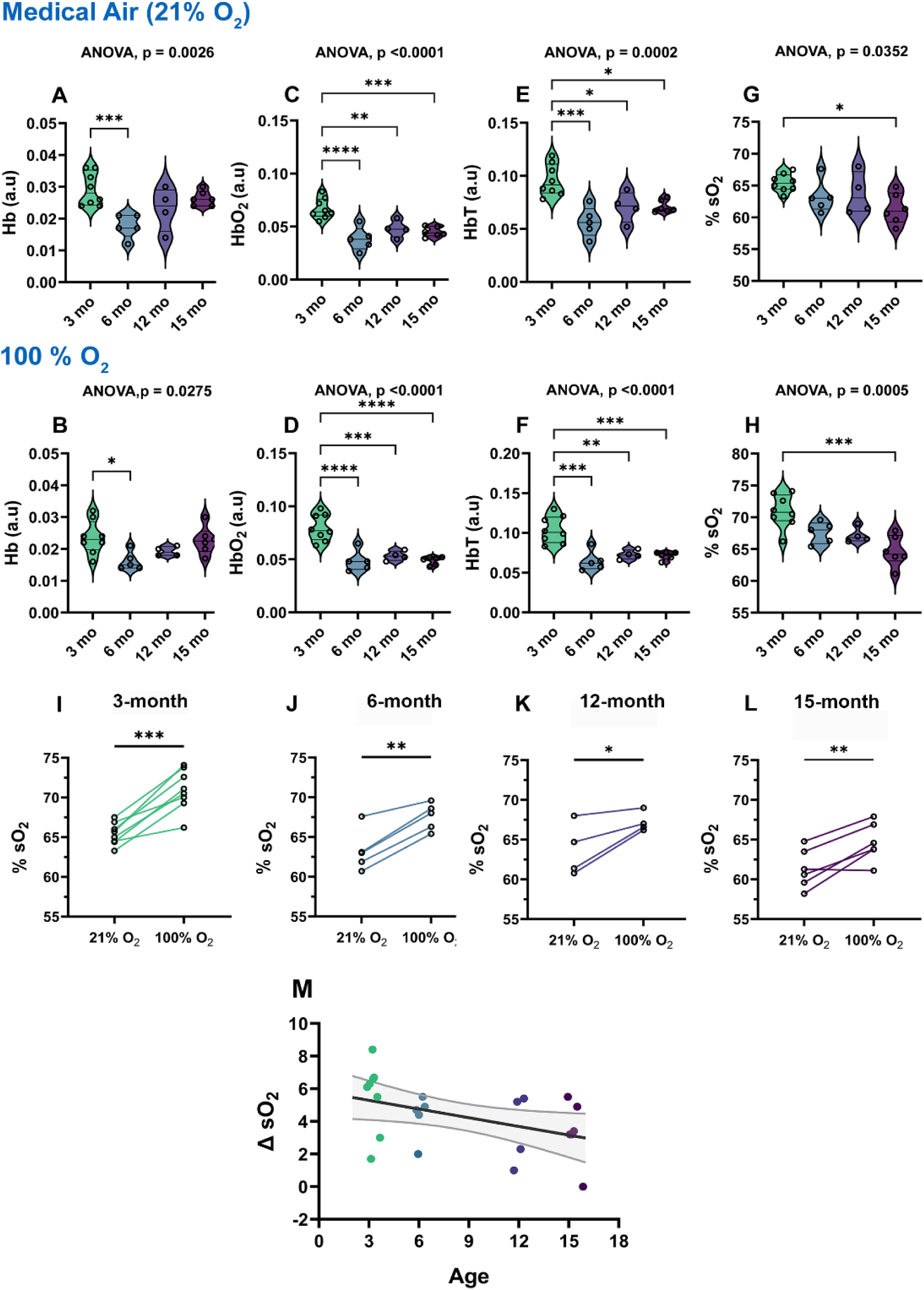
Lower cerebral HbO₂, HbT and sO₂^MSOT^ and a reduced oxygen-challenge response in older mice. (A-H) Whole-slice Hb, HbO₂, HbT and sO₂^MSO^T in mice aged 3, 6, 12 and 15 months (n = 8, 5, 4 and 6, respectively) under 21% O₂ (A, C, E, G) and 100% O₂ (B, D, F, H). The p-value for the effect of age (one-way ANOVA) is shown above each panel; asterisks indicate Dunnett’s multiple comparisons test against the 3-month group. (I–L) Whole-slice sO₂^MSOT^ under 21% and 100% O₂ in individual mice at 3 (I), 6 (J), 12 (K) and 15 (L) months, with lines connecting paired measurements from the same animal (two-tailed paired t-test). (M) Oxygen-challenge response, ΔsO₂ plotted against age, with the linear regression fit and its 95% confidence band. *p < 0.05, **p < 0.01, ***p < 0.001, ****p < 0.0001. a.u., arbitrary units

Hb, HbO_2_ and HbT differed between the age groups independent of inspired air (one-way ANOVA: HbO_2_, p<0.0001 under both 21% and 100% O_2_; HbT, p=0.0002 and p<0.0001 and Hb, p=0.003 and p=0.03, under 21% and 100% O_2_ respectively; **Fig. 4A-F**). Compared with 3-month-old mice, HbO₂ and HbT were lower in all older groups (Dunnett’s multiple comparisons test). Under 21% O₂, HbO₂ and HbT in 15-month-old mice were 33% (Hedges’ g=2.6) and 26% (g=1.9) lower, respectively, than in 3-month-old mice. Although Hb differed between groups, the difference was only significant between 3-month and 6-month but did not vary significantly between 3 and 15 months. sO₂^MSOT^ was calculated pixel-wise as (Equation. 1) and averaged within the ROI and reported as a percentage. %sO₂^MSOT^ differed between the age groups (21% O₂, p=0.035; 100% O₂, p=0.0005; **Fig. 4G, H**). The difference was significant between 3 and 15 months, with intermediate values at 6 and 12 months. At 15 months, sO₂^MSOT^ was 4.1 percentage points lower under 21% O₂ (g=2.0) and 6.3 percentage points lower under 100% O₂ (g=2.3). Because sO₂^MSOT^ reflects the balance between oxygen delivery and consumption in the sampled tissue, lower values in aged mice are consistent with reduced oxygen delivery, increased oxygen extraction, or both.

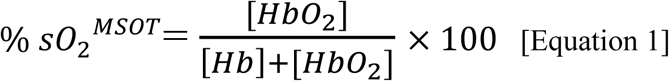

### 3.4 The oxygen challenge response decreases modestly with age

To probe the response of the cerebral vasculature to hyperoxia (47, 48), the fraction of inspired oxygen was increased from 21% to 100% O₂, and the resulting change in %sO₂^MSOT^ within the whole-slice ROI was quantified in mice aged 3, 6, 12 and 15 months (**Fig. 4I-L**).

In every age group, %sO₂^MSOT^ increased from 21% to 100% O₂ (paired t-test: 3 months, p=0.0002; 6 months, p=0.002; 12 months, p=0.04; 15 months, p=0.009; **Fig. 4I-L**). Under 100% O₂, %sO₂^MSOT^ remained lower in 15-month-old than in 3-month-old mice (**Fig. 4H**), so the age-group difference persisted at maximal inspired oxygen. The magnitude of the oxygen-challenge response, ΔsO₂^MSOT^=%sO₂^MSOT^ (100% O₂)-%sO₂^MSOT^ (21% O₂), decreased with age (**Fig. 4M**). Linear regression of ΔsO₂^MSOT^ gave a negative slope of 0.18% per month (95% CI −0.34 to −0.01; p=0.034; n=23), corresponding to an estimated reduction of 2.2% (95% CI 0.2-4.1) between 3 and 15 months.

Taken together, %sO₂^MSOT^ increased in response to hyperoxia at every age examined but was persistently lower in 15-month-old than in 3-month-old mice under both 21% and 100% O₂. The magnitude of the oxygen-challenge response (ΔsO₂^MSOT^) decreased modestly with age, although differences between age groups were not significant, reflecting the small group sizes and inter-animal variability.

### 3.5 Microvascular rarefaction, loss of GLUT1 and reduced microglial ramification in aged brain

To relate imaging findings to tissue structure, brain sections from imaged 3-month (n=6) and 15-month-old mice (n=7) were immunostained for the endothelial marker CD31, the microglial marker IBA1 and the endothelial glucose transporter (GLUT1, n=2 from the two age groups). (**Fig. 5**). In 3-month-old mice, CD31 displayed a dense, interconnected microvascular network whereas vessels in 15-month-old mice appeared sparser and more fragmented (**Fig. 5A, B**). GLUT1 likewise labeled a continuous network in 3-month-old brains, whereas in 15-month-old brains GLUT1⁺ labeling was discontinuous and confined to short vessel segments (**Fig. 5C, D**; n=2 mice per age, assessed qualitatively). In both age groups, IBA1 microglia had small somata and ramified processes typical of non-activated microglia, with no amoeboid forms; however, microglia in 15-month-old mice had fewer and less elaborate processes (**Fig. 5E, F**).

**Figure 5.**
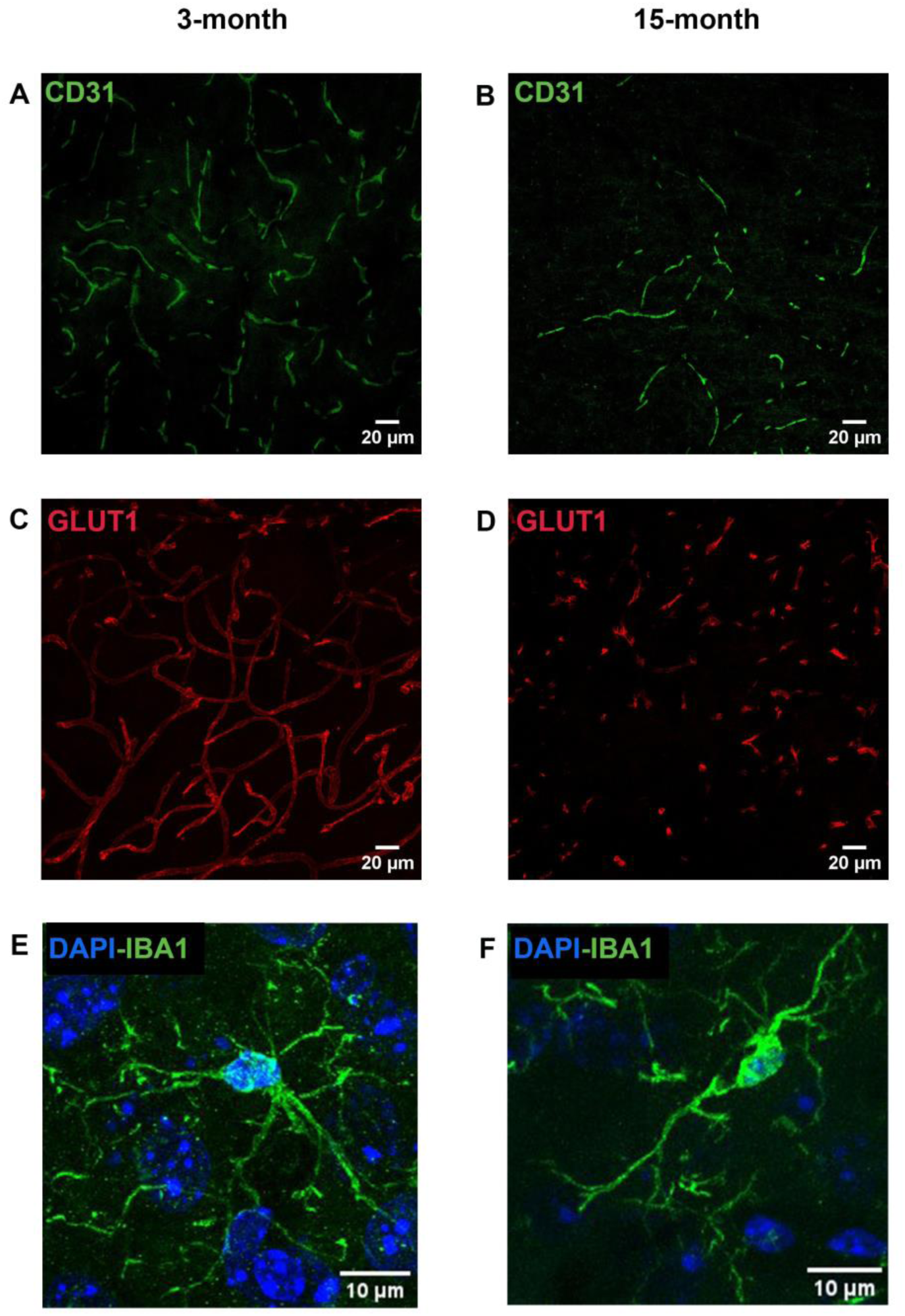
Sparser microvessels, reduced GLUT1⁺, and less ramified microglia in the aged cortex. Representative confocal images (maximum-intensity projections of 50 µm z-stacks) from the cortex at approximately Bregma 0 to −1 mm. (A, B) CD31⁺ microvessels in 3-month-old and 15-month-old mice, (40x magnification). (C, D) GLUT1 immunostaining in (C) 3-month-old and (D) 15-month-old mice. (E, F) Iba1⁺ microglia (green) with DAPI nuclear counterstain (blue) in (E) 3-month-old and (F) 15-month-old mice, (60x magnification). Images were acquired and displayed with identical settings for both age groups. Scale bars, 20 µm (A-D) and 10 µm (E, F).

### 3.6 MSOT blood-volume readouts track CD31^+^ vascular density across and within age-groups

Animal-level quantification confirmed these observations. CD31⁺ vascular density, averaged over 10 fields (184 × 184 µm²) per mouse, was 56% lower in 15-month-old than in 3-month-old mice (2.38 ± 0.45% vs 1.04 ± 0.54% area, p=0.0003), and vascular length density was reduced by a similar proportion (−55%; p=0.0007) (**Fig. 6C, D**). Skeleton analysis of IBA1⁺ microglia (10-15 cells per mouse, averaged per animal) showed fewer branches (−26%; p=0.04), junctions (−29%; p=0.04) and lower branch length (−16%; p=0.02) per cell in 15-month-old mice compared to 3-month-old mice (**Fig. 6A, B**, **Fig. S6)**.

**Figure 6.**
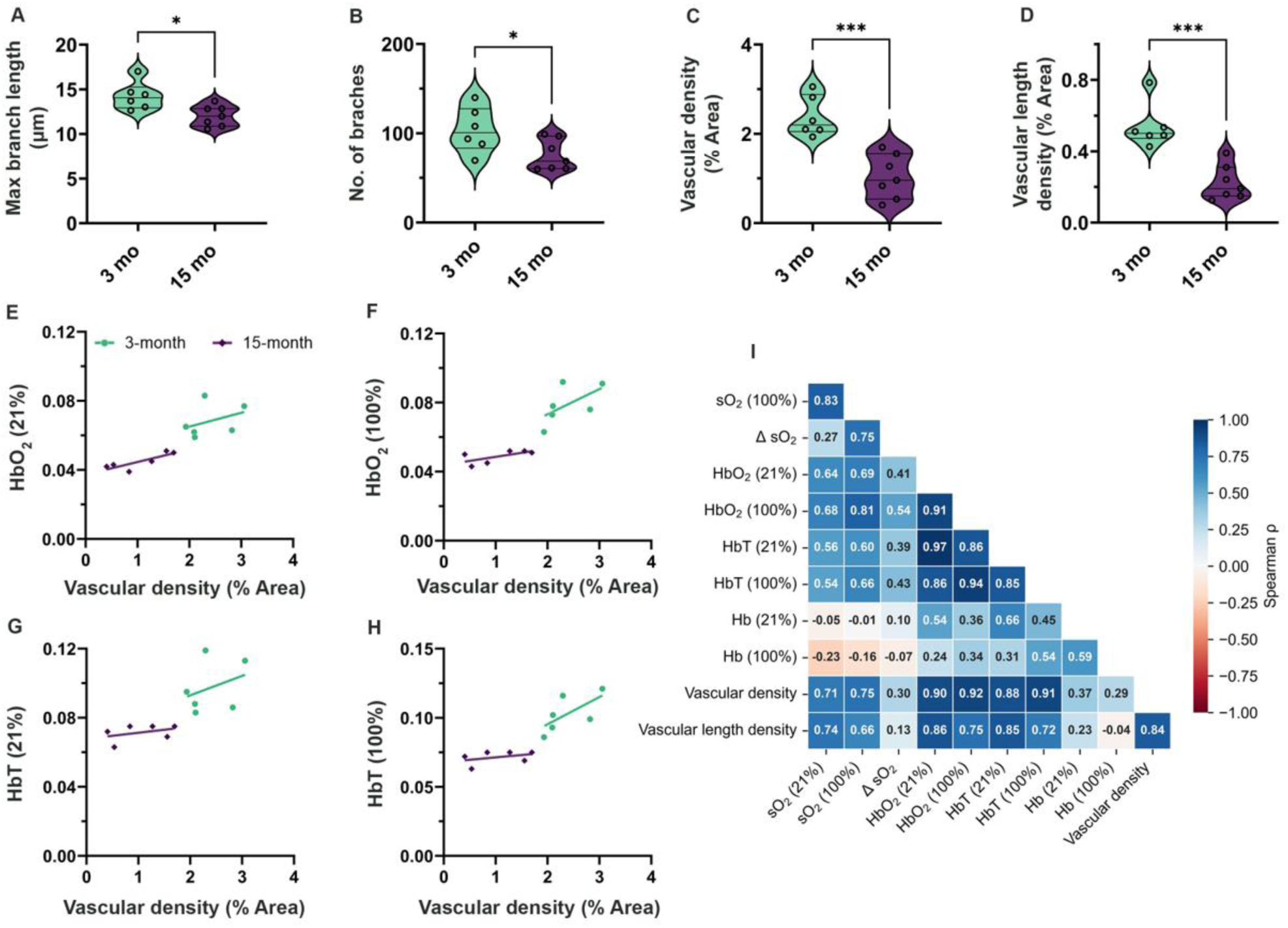
Microglial morphology and microvascular density in 3- and 15-month-old mice, and their relationship to MSOT readouts. (A, B) Skeleton analysis of Iba1⁺ microglia: (A) maximum branch length and (B) number of branches per cell. (C, D) CD31⁺ vascular density (C) and vessel length density (D) in the cortex. Each point represents one mouse. Groups were compared with unpaired two-tailed t-tests with Welch’s correction. (E-H) Relationship between CD31⁺ vascular density and (E) HbO₂ under 21% O₂, (F) HbO₂ under 100% O₂, (G) HbT under 21% O₂ and (H) HbT under 100% O₂, with linear regression lines fitted separately for each age group (n = 12). (I) Spearman correlation matrix (*ρ*) of MSOT readouts and histological measures, pooled across both age groups (n = 12). Additional microglial measures are shown in Supplementary Fig. S6. *p < 0.05, ***p < 0.001

We next compared the MSOT readouts with vascular length density in the same animals (n=12; **Fig. 6E-I**). Across all animals, CD31⁺ vascular density correlated strongly with HbO₂ and HbT (Spearman ρ=0.90-0.92, p<0.001) and with sO₂^MSOT^ (ρ=0.71–0.75, p≤0.009), but not with ΔsO₂^MSOT^ (ρ=0.30, p=0.34) or Hb (ρ=0.29-0.37, p≥0.24) (**Fig. 6I**). Since the pooled correlations partly reflect the age-group differences for both vascular density and MSOT readouts, associations were also evaluated within each age-group. HbO₂ and HbT showed positive associations with vascular density at both ages (3 months, r=0.34-0.66; 15 months, r =0.39-0.87), with similar slopes in the two age groups (comparison of slopes, p≥0.19, **Fig 6E-H**). After adjustment for age (ANCOVA), these associations remained positive (partial r= 0.44-0.52) but did not reach statistical significance at this sample size (p=0.10-0.18) reflecting the strong collinearity between age and vascular density at this sample size. In the same models, HbO₂ under 100% O₂ remained lower in 15-month-old mice after adjustment for vascular density (age effect, p=0.04); however, this age effect was not significant for the other readouts (p≥0.09). Microglial branching correlated only moderately with MSOT readouts across all animals (*ρ* ≤0.58) and was not associated with any readout after adjustment for age, indicating that the age-related changes in microglial morphology occurred in parallel with, rather than in proportion to the hemodynamic readouts.

## 4. Discussion

In this study, OE-MSOT with NNMB reconstruction detected age-group differences in cerebral hemodynamics in healthy female BALB/c mice, and these differences were mirrored by histological microvascular rarefaction in the same animals. Three main findings can be highlighted from this study. First, MSOT-derived HbO₂, HbT and %sO₂^MSOT^ were lower in older mice under both normoxia and hyperoxia, with the largest differences in the blood-volume (HbT) readouts. Second, these readouts correlated strongly with CD31⁺ vascular density strongly across animals, and with a consistent direction within each age group. Third, the oxygen-challenge response (ΔsO₂^MSOT^) declined modestly with age. Together, these results indicate that MSOT blood-volume readouts reflect the underlying microvasculature and can be used to follow cerebrovascular aging noninvasively.

Among the MSOT readouts, HbT reflects the intravascular hemoglobin content in tissue and is widely used as a surrogate for cerebral blood volume (CBV)(54, 55), whereas sO₂^MSOT^ reflects the local balance between delivery and consumption of oxygen and is expected to be weighted toward venous oxygen saturation, because most of the CBV is venous and capillary(29). The lower HbT in aged mice concurs with the reduction in CD31⁺ vascular density, in agreement with previous reports of age-related capillary rarefaction, reduced vascular branching and pericyte loss in the rodent brain(1, 56, 57). The two measures were also strongly correlated across animals in our study. Because both differ with age, however, a key question that arises is whether this correlation simply reflects their shared dependence on age.

For each age group, HbO₂ and HbT remained positively associated with vascular density, with similar slopes at both ages, but these associations did not reach significance after adjustment for age. The lack of significance, though, is assumed to be largely a consequence of design: age and vascular density were strongly collinear, leaving little independent variation to test with six animals per group. The data are therefore consistent with MSOT readouts tracking vascular density within as well as between ages, but larger cohorts will be needed to confirm this. Notably, under hyperoxia, HbO₂ remained lower in aged mice even after accounting for vascular density, suggesting that factors beyond vessel number, such as oxygen delivery per vessel, may also contribute. Lower baseline %sO₂^MSOT^ in aged mice is likewise consistent with reduced oxygen delivery, increased oxygen extraction, or both.

Hyperoxia increased %sO₂^MSOT^ at every age, and the magnitude of this response declined modestly with age. Hyperoxia causes mild cerebral vasoconstriction(31, 58), and the resulting rise in tissue oxygenation reflects a combined effect of increased arterial oxygen content and this vasoconstriction on oxygen delivery. The simplest explanation for the smaller response in aged brains would be fewer vessels and lower blood volume. However, because sO₂^MSOT^ is a ratio, lower blood volume alone would not be expected to reduce the response; and indeed, ΔsO₂^MSOT^ was only weakly and not significantly correlated with vascular density (*ρ* =0.30, p =0.34). The smaller response in aged mice may instead reflect reduced oxygen-carrying capacity of the blood(59), increased baseline oxygen-extraction(6) or age-related differences in hyperoxic vasoconstriction. Similarly, ΔsO₂^MSOT^ should not be equated with cerebrovascular reactivity, which is assessed with vasodilatory stimuli such as hypercapnia and is reduced early in aging and age-related disease(60, 61). Combining hyperoxic and hypercapnic challenges with blood-flow measurements in future studies would allow both aspects to be assessed in the same animals. A useful comparison comes from oxygen-enhanced optoacoustic studies in tumors, where dynamic metrics such as ΔsO₂ were more repeatable than static measures and correlated more closely with histological markers of vascular function and hypoxia (36, 43). In the healthy brain, we observed the opposite pattern: static blood-volume readouts were more closely related to histology, whereas ΔsO₂^MSOT^ changed only modestly with age and varied widely between animals, limiting its sensitivity. This likely reflects the much narrower dynamic range of oxygenation in well-perfused brain tissue than in hypoxic tumors.

Two recent non-photoacoustic studies provide important insights: Bennett et al. reported a reduction in vascular length density in 18-month-old C57BL/6 mice, concentrated in deep cortex, hippocampus and basal forebrain, together with reduced oxygen delivery during 100% O₂ breathing(59), consistent with our smaller oxygen-challenge response. However, the larger CD31⁺ reduction we observed compared to the aforementioned study likely reflects differences in strain, region and the method (CD31⁺ area fraction in sections vs 3D whole-brain vascular tracing). A longitudinal MRI study found preserved cerebral blood flow but increased oxygen consumption up to 20 months(6), supporting increased oxygen extraction as a contributor to lower sO₂^MSOT^. To our knowledge, the present work is the first to relate photoacoustic readouts of the aging brain to microvascular histology in the same animals.

An unexpected finding of this study was how strongly the reconstruction algorithm shaped the results. Standard BP compressed the dynamic range of sO₂^MSOT^, so that the hyperoxic response appeared roughly three-fold smaller than with NNMB and the age-group difference was diminished, consistent with negative-value artifacts distorting the spectral unmixing(42). Although the age difference was present with both methods, only NNMB resolved it with statistical confidence. A larger effect size does not by itself indicate greater accuracy. Rather NNMB was preferred because it enforces a physically meaningful non-negativity constraint, and because non-negative constrained model-based inversion has been shown to improve image accuracy and the reliability of spectral unmixing compared with unconstrained reconstruction(62). In practice, subtle physiological effects may be detected with one reconstruction and missed with another, so reconstruction methods and settings should be reported in detail in photoacoustic neuroimaging studies.

Microglia in 15-month-old mice retained a ramified morphology without amoeboid features, but had fewer branches, reduced maximum branch length and junctions than in the young mice. This simplification of the process arbor is consistent with age-related microglial remodeling described in the healthy aging brain, as distinct from the reactive transformation seen in neuroinflammation or neurodegeneration(23, 63). Microglial branching was not associated with the MSOT readouts after adjustment for age, indicating that these changes occur in parallel rather than in proportion to the hemodynamic differences.

Our GLUT1 staining points toward a connection between vascular and metabolic aging. As the principal glucose transporter of brain endothelium, GLUT1 governs glucose entry across the blood-brain barrier(64), with reduced blood-brain barrier glucose transport being reported in aged rodents(65), including an early decline in female mice(66). Loss of endothelial GLUT1 has also been shown to aggravate neurovascular dysfunction in a model of Alzheimer’s disease (67). The reduced GLUT1⁺ labeling in aged brains is consistent with these reports and suggests that, alongside fewer vessels, the remaining microvasculature may have reduced glucose-transport capacity. Because GLUT1 was assessed in only two animals per age group, these observations require confirmation and co-localization with CD31^+^ in larger cohorts. If confirmed, they would also be relevant to glucose-targeted imaging probes, whose delivery to the aged brain may be reduced.

From a translational perspective, OE-MSOT is complementary to many established clinical methods. PET provides absolute measures of cerebral blood flow and oxygen metabolism using ¹⁵O-labeled tracers (25, 26) but is difficult to apply repeatedly in mice because of tracer production, radiation dose and limited spatial resolution. MSOT provides relative measures of hemoglobin content and oxygenation without ionizing radiation or exogenous tracers, and can be used for longitudinal assessment. It is therefore well suited to preclinical studies of cerebrovascular aging and of interventions designed to slow it, such as exercise, dietary restriction or senolytic treatment, with less frequent PET or MRI used for calibration. In adult humans, the skull remains the main barrier to transcranial photoacoustic imaging. Translation is therefore more immediately feasible where this barrier is reduced or absent, such as imaging the neonatal brain or the exposed cortex during neurosurgery, or by measuring microvascular oxygenation in peripheral tissues as an accessible proxy for systemic vascular aging(68).

A few limitations should be considered while evaluating these results from our study. First, the MSOT readouts are relative rather than absolute. Fluence correction relies on a generic tissue model, and although sO₂^MSOT^, as a ratio, cancels global signal attenuation, it remains sensitive to wavelength-dependent attenuation. Second, age-related changes in skull thickness and composition may alter light delivery and acoustic transmission. In our results, scalp signals did not differ between age groups, but this does not completely exclude skull effects on the brain signal. The within-age association with vascular density and the within-scan ΔsO₂^MSOT^ are less susceptible to such effects, but direct assessment, for example by micro-CT of skull thickness, could be useful. Finally, imaging was performed under isoflurane, which is vasodilatory. Although a fixed gas order (21% followed by 100% O₂) along with a set anesthetic dose and timing were standardized across animals, yet anesthesia may still have modified the hemodynamic responses to some extent. Despite these limitations, the consistency between imaging and histology across animals supports the use of OE-MSOT readouts as noninvasive markers of cerebrovascular aging.

## Conclusion

In summary, OE-MSOT with NNMB reconstruction detected lower blood-volume and oxygenation readouts and a smaller oxygen-challenge response in the aging mouse brain, and the blood-volume readouts tracked histological microvascular rarefaction in the same animals. These results support OE-MSOT as a noninvasive, radiation-free tool for longitudinal studies of cerebrovascular aging and of interventions that target it.

## Data availability

Data will be made available on request.

## Acknowledgments

The authors would like to acknowledge the Central Animal Facility, IISc, for their support with animal work. The authors would like to also acknowledge the J N Tata MRI Centre, IISc for their support.

## Author contribution

Roshni Parameswaran: Methodology, investigation, formal analysis, reviewing. B Thamaraiselvi: Methodology, formal analysis, writing and reviewing. Ankita Mandal: Methodology, review and writing, formal analysis. Thomas Sardella: Data curation, analysis, reviewing. Sanhita Sinharay: Conceptualization, Supervision, validation, writing original draft, investigation, funding acquisition.

## Ethics declarations

This study was performed in line with the Institutional Animal Ethics Committee, Indian Institute of Science (protocol no. IISc CAF/ethics/099/2024).

