## Supplementary figures for "Oxygen-enhanced photoacoustic tomography of cerebral hemodynamics in the aging mouse brain, validated against microvascular and microglial histology"

**Figure S1. $HbO_2$ and Hb during the oxygen challenge.** Representative dynamic acquisition in the same 3-month-old mouse as in **Fig. 2C**, showing (A) $HbO_2$ and (B) Hb within the whole-slice ROI during the switch from 21% to 100% $O_2$ (arrow). $HbO_2$ increased and Hb decreased reciprocally after the gas switch, accounting for the rise in $sO_2^{MSOT}$ while HbT remained stable (**Fig. 2C**)

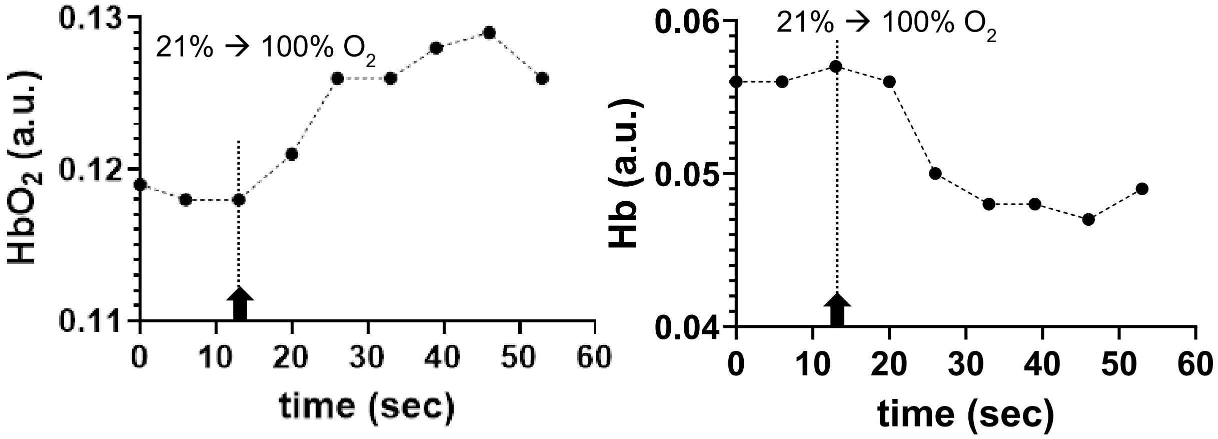


**Figure S2**: (A, B) Qualitative images of the whole brain slice without and with the scalp ROI on a 3-mo-old and 15-mo -old mouse brain, (C, D) optoacoustic intensity at 800 nm and HbT (a.u.) in the scalp ROI for mice in both age-groups.

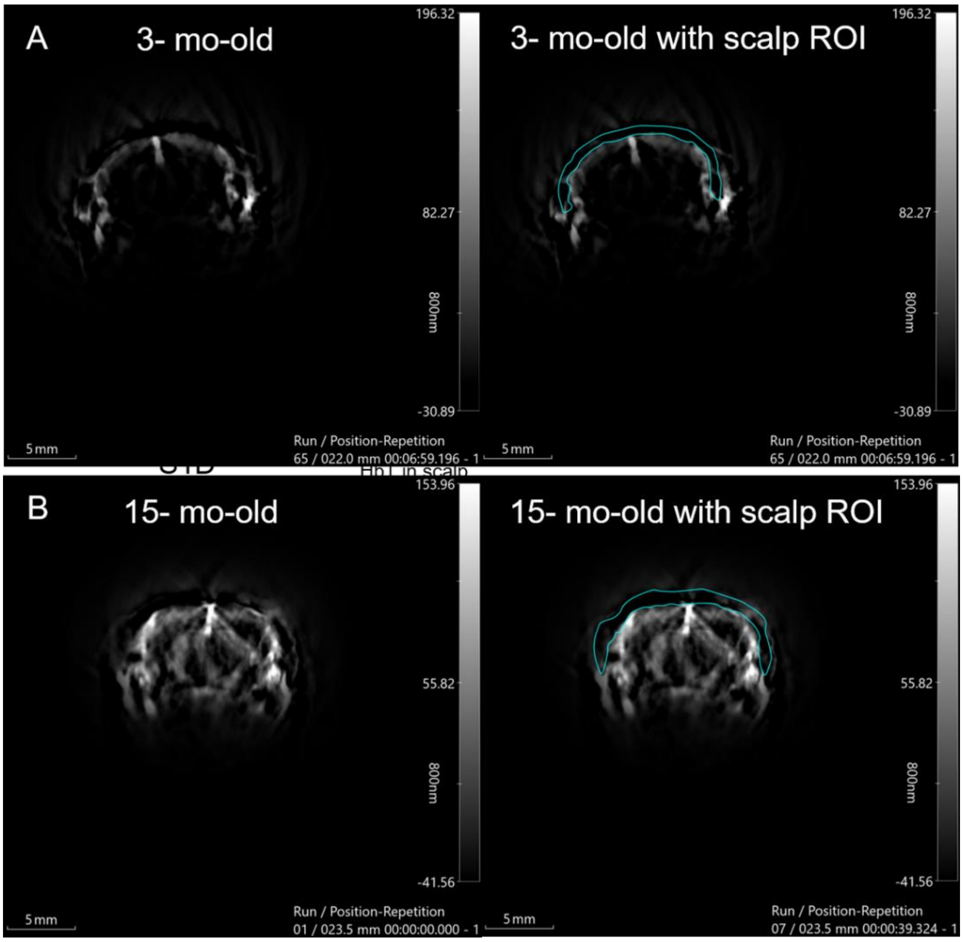

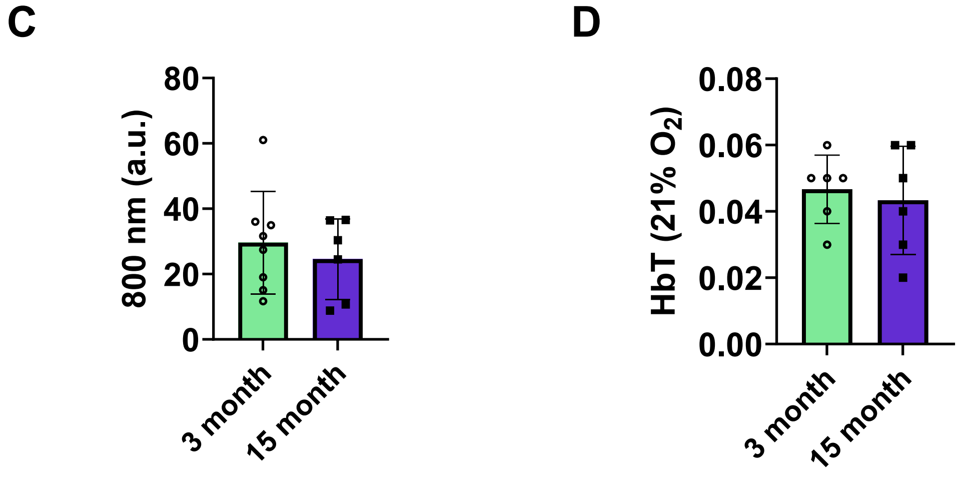


**Figure S3**: **Regional $sO_2^{MSOT}$ within the imaging plane.** (A) Regions of interest in a representative 3-month-old mouse: whole slice (WS), cortex (C), striatum (S), midline vessels and lateral surface vessels. (B) $sO_2^{MSOT}$ in each region under 21% and 100% $O_2$ ($n = 1$). Cortical and striatal $sO_2^{MSOT}$ were similar to the whole-slice value, and all regions increased under hyperoxia. Scale bar, 5 mm

**A**

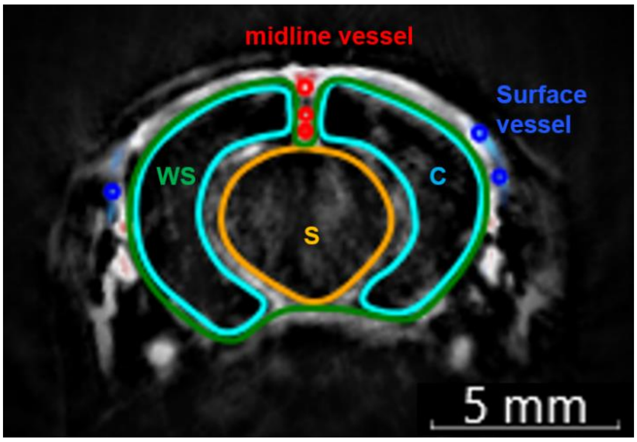


**B**

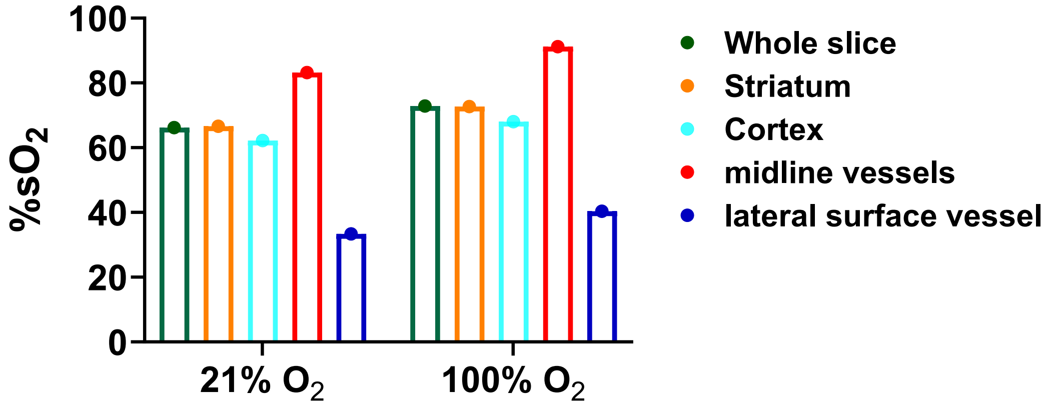

**Supplementary Fig. 4: Immunofluorescence data of mice not included in the main figures:** Cortical microvascular rarefaction and altered microglial morphology with aging. (A-D) Confocal images of cortical vessels stained for CD31 (A, B) and GLUT1 (C, D) in 3-month-old (A, C) and 15-month-old (B, D) mice. Scale bars, 20 µm. (E, F) Iba1 positive microglia (green) with DAPI nuclear counterstain (blue) in the cortex of 3-month-old (E) and 15-month-old (F) mice. Scale bars, 10 µm.

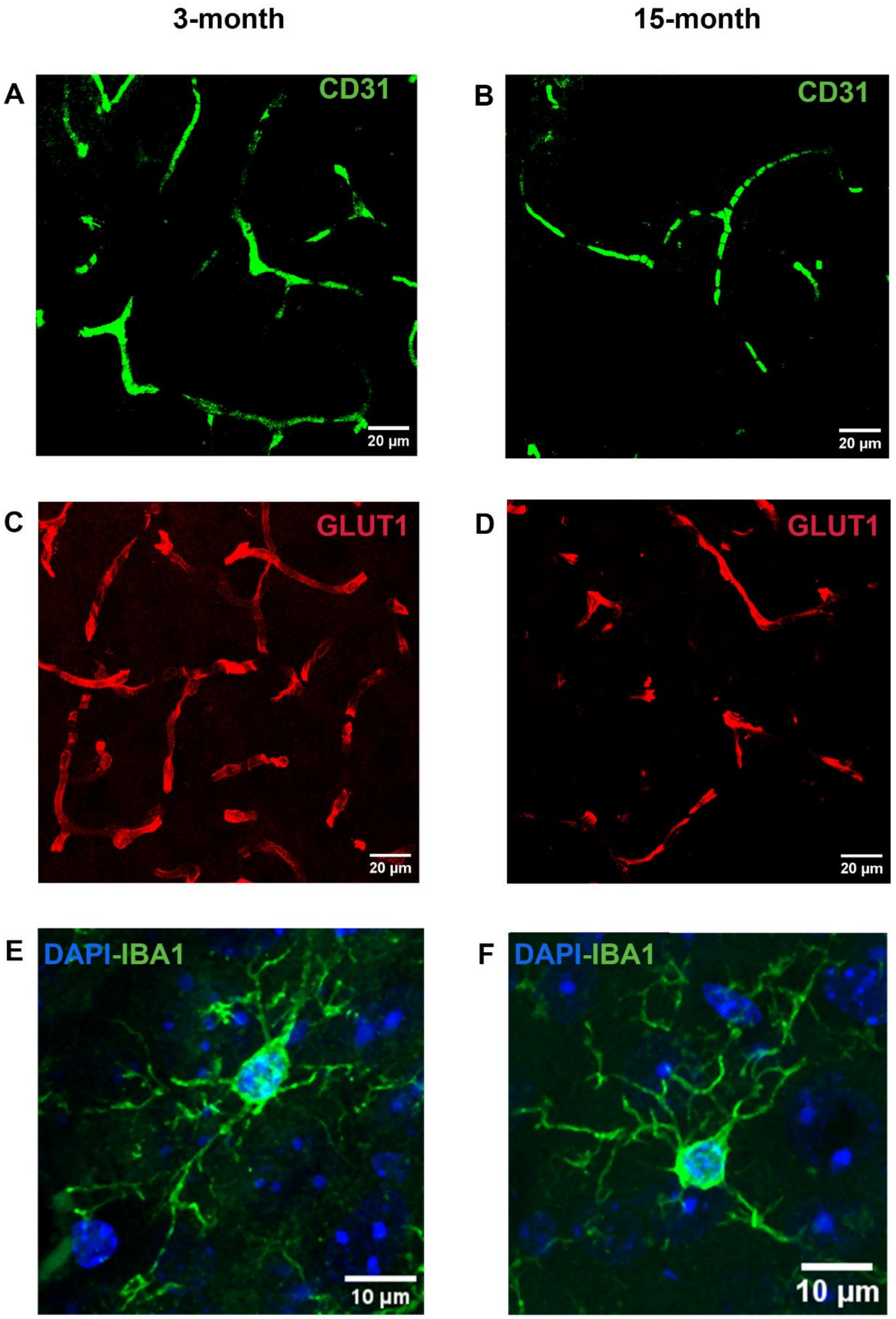

**Supplementary Fig. 5: Representative coronal brain section near bregma used for immunofluorescence analysis (IFA).** Brightfield (transmitted-light) tile-scan overview of a 50 µm coronal section from a 3-month-old mouse, acquired on a Leica TCS SP8 confocal microscope with a 20×/0.40 dry objective and merged in LAS X software**.**

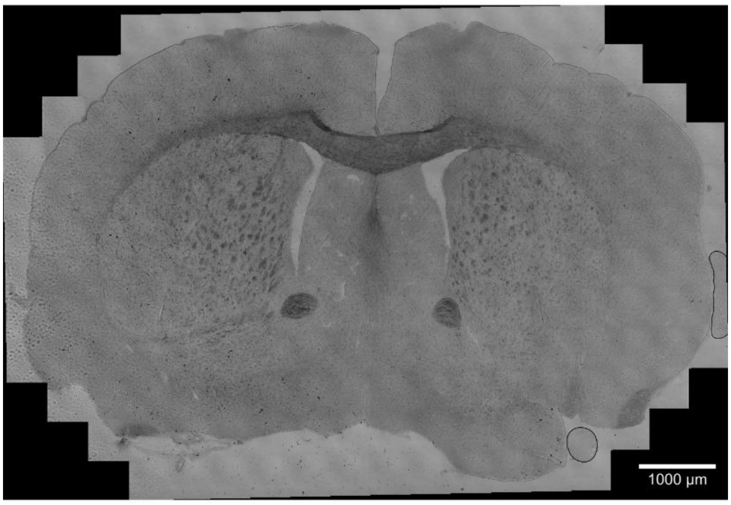


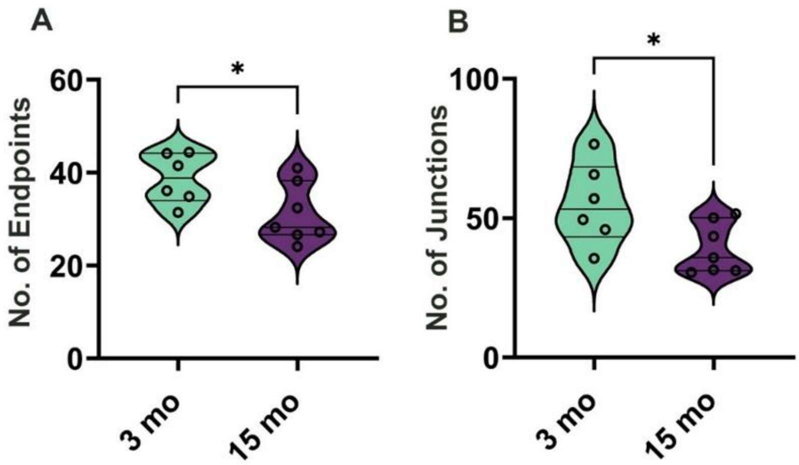


**Figure S6. Additional microglial morphometry in 3- and 15-month-old mice.** Number of endpoints (A) and junctions (B) per Iba1$^{+}$ microglial cell, obtained by skeleton analysis. Each point represents one mouse (mean of 10-12 cells per animal); 3 months, n = 6; 15 months, n = 7. Both measures were lower in 15-month-old mice (endpoints, −18%; junctions, −29%; unpaired two-tailed t-test with Welch's correction, *$p < 0.05$)
